# Beyond dairy: Identification of dental enamel proteins in ancient human dental calculus

**DOI:** 10.64898/2026.03.21.713223

**Authors:** Adriana Leite, Frido Welker, Ricardo Miguel Godinho, Rosalind Emma Gillis, Viridiana Villa-Islas, Zandra Fagernäs

**Affiliations:** ICArEHB – The Interdisciplinary Center for Archaeology and Evolution of Human Behaviour, Universidade do Algarve, Faro, Portugal; Globe Institute, University of Copenhagen, Øster Voldgade 5-7, Copenhagen, Denmark; Referat Naturwissenschaften, DAI, Berlin, Germany

**Keywords:** Dental Calculus, Palaeoproteomics, Enamel Proteome, Amelogenin, Sex Estimation

## Abstract

Ancient human dental calculus is one of the richest archives of archaeological biomolecular information, providing direct evidence of diet, oral health, and the oral microbiome. Proteomic analyses of this biological matrix have so far focused mainly on oral microbes and dietary proteins, with milk proteins such as beta-lactoglobulin (BLG) providing the largest corpus of proteomic evidence. Despite the close relation between the various stages of dental calculus formation and mineralization with the dental enamel surface, proteins from the dental enamel matrix have not previously been reported outside of dental enamel tissue. Here we reanalysed 498 ancient dental calculus proteomes from 14 published studies (n=434 individuals) reporting the presence of BLG, spanning from the Neolithic to the Victorian Era and applying different protein extraction protocols (FASP, GASP, SP3 and in-solution digestion). Dental enamel matrix proteins were identified in ten studies (n=37 individuals), with amelogenin being the most frequently detected. Enamel peptides occurred more often in studies that applied SP3, although amelogenin was successfully identified through both SP3 and FASP. Structural proteins, including enamelin, ameloblastin, and MMP20, were also identified. The detection of AMELX and AMELY peptide sequences provided new insights into cases where the sex was previously undetermined. These findings establish dental enamel proteins as a new category of biomolecules detected in dental calculus, broadening its application beyond diet and microbiome studies to possible sex estimation.

**Highlights:**

- Dental calculus entraps oral microbes along with endogenous and exogenous particles during formation and mineralization
- We conduct reanalysis of 14 published ancient dental calculus studies (n = 434 individuals) spanning the Neolithic to Victorian Era
- Dental enamel proteins AMELX, AMELY, AMBN, COL17A1, ENAM and MMP20 are identified in ancient human dental calculus
- Amelogenin was the most frequently detected enamel protein
- We expand dental calculus palaeoproteomics beyond diet and oral microbiome to potentially include sex estimation

## 1. Introduction

Dental calculus (tooth tartar) is a complex calcified plaque matrix that accumulates throughout an individual’s lifetime and is considered a long-term reservoir of the oral microbiome (Forshaw, 2022; Kinaston et al., 2019; Warinner et al., 2015). Although long overlooked in the past, archaeological dental calculus is now well-established as a powerful source of information on oral health, disease and dietary practices (Mackie et al., 2017a; Warinner et al., 2015). Over the past decade, one of the major research *foci* has been the reconstruction of specific food components through the extraction of dietary proteins preserved in calculus, yielding direct evidence of cereal and plant consumption (e.g. oats, peas, wheat, sesame, soybean) (Hendy et al., 2018; Scott et al., 2021), and animal-derived products, particularly dairy, throughout prehistory (e.g., Bleasdale et al., 2021; Charlton et al., 2019; Jeong et al., 2018; Scott et al., 2022; Tang et al., 2023; Ventresca Miller et al., 2023, 2022; Warinner et al., 2014; Wilkin et al., 2021, 2020). Notably, the largest corpus of proteomic evidence generated from calculus relates to dairy consumption, as milk proteins are among the best-preserved dietary markers (Fagernäs and Warinner, 2023). Specifically, beta-lactoglobulin (BLG) has received particular attention as it is the most frequently recovered dietary protein, owing to its stability and resistance to degradation, which may be related to its ability to bind to mineral surfaces (Fonseca et al., 2022; Warinner et al., 2022).

The processes that lead to calculus formation take place in the oral cavity, which harbors a highly diverse microbial community (Hillson, 2005; Warinner, 2016). Soon after tooth eruption, salivary proteins present in the mouth selectively adsorb onto the enamel surface to form the acquired enamel pellicle (AEP) (Jin and Yip, 2002; Lendenmann et al., 2000). Among the first to adsorb are salivary proteins with high affinity to hydroxyapatite – including statherin, histatins, proline-rich proteins, cystatins, amylase and lysozyme – which can undergo exchange reactions with the calcium and phosphate ions of the enamel (Ferrari et al., 2025; Hannig and Joiner, 2005). The AEP plays an important protective role by reducing abrasion through the formation of a barrier between the enamel surface and the oral cavity, while also maintaining tooth integrity by modulating the demineralization– remineralization process and thereby slowing the rate of mineral dissolution (Hannig and Joiner, 2005; Lendenmann et al., 2000; Ventura et al., 2017). This is followed by the adherence of oral microbes (primary or early colonisers) to the pellicle-coated enamel surface, and the subsequent attachment of secondary or later colonisers through co-aggregation, resulting in the development of a dental plaque biofilm (Ferrari et al., 2025; Jin and Yip, 2002; Marsh and Bradshaw, 1995). Mineralization of the dental biofilm occurs via the deposition of calcium and phosphate from saliva and gingival crevicular fluid, leading to the formation of dental calculus (Mandel, 1969; White, 1997). Through repeated cycles of plaque formation and mineralization, calculus accumulates incrementally, entrapping microparticles and biomolecules from food, plant and animal fibers, and environmental debris, eventually forming a cement-like deposit (Fagernäs and Warinner, 2023; Kinaston et al., 2019; White, 1997).

Despite the close interaction between the tooth surface and dental calculus during the various stages of its formation, proteins from the enamel matrix have not previously been reported outside of the enamel tissue. The small proteome of enamel consists primarily of structural proteins including amelogenin, ameloblastin, enamelin, amelotin, odontogenic ameloblast-associated protein and tuftelin, together with enamel proteases secreted during amelogenesis, including matrix metalloproteinase-20 and kallikrein-4 (Cappellini et al., 2019; Warinner et al., 2022). Among these, amelogenin is the most abundant enamel protein and occurs in two sexually distinct forms, AMELX and AMELY, allowing the estimation of genetic sex in archaeological specimens (Castiblanco et al., 2015; Koenig et al., 2024). The presence of both AMELX and AMELY indicates a male individual, whereas the detection of AMELX alone is inconclusive as it may reflect either a female individual or a low signal male sample with no detectable AMELY peptides (Parker et al., 2019; Welker et al., 2025).

Here, we explored the preservation of proteins from dental enamel in ancient human dental calculus samples from 434 individuals from previously published studies reporting the detection of BLG. These studies represent high-quality palaeoproteomic data currently available for ancient calculus, spanning a wide chronological range from the Neolithic to Victorian Era and a broad geographical distribution. The successful identification of BLG and other dietary proteins indicates good biomolecular preservation, making them particularly suitable for re-analysis. While the original studies focused mainly on a specific subsistence practice of past communities – dairying – a significant portion of the proteomic data extracted from these archaeological specimens remains unexplored. As dental calculus sampling and proteomic examination is destructive, the analysis of existing datasets provides an excellent opportunity to address new research questions without the need of sampling new archaeological material. Drawing from this large ancient proteomic dataset, we compared enamel proteome preservation across extraction methods (filter-aided sample preparation (FASP), gel-aided sample preparation (GASP), SP3, and in-solution digestion), and presence or absence of dairy proteins. Where available, sex estimation from amelogenin was compared with osteological and genetic sex determinations. Together, our analysis shows that enamel proteins can survive in dental calculus across diverse archaeological contexts and establish them as a new category of biomolecules preserved in this complex matrix.

## 2. Materials and methods

### 2.1 Literature search

A comprehensive search of peer-reviewed studies that had identified milk proteins, specifically BLG, in ancient human dental calculus, was conducted in May and June 2025 using the search engine Google Scholar. The search terms were the following: “dental calculus”, “ancient human dental calculus”, “palaeoproteomics”, “milk consumption”, “beta-lactoglobulin” and “BLG”. From our search, we identified 14 studies published between 2014 and 2023 that met our inclusion criteria. Studies focusing solely on the optimization and validation of protein extraction methods from dental calculus were excluded.

For the selected studies, all the available metadata were compiled (Supplementary Information 1), including archaeological and extraction laboratory identifiers, archaeological period or cultural designation, sample size, osteological sex estimation, genetic sex identification, presence of dairy proteins, and extraction methods. In the case of Jersie-Christensen et al. (2018), the archaeological IDs corresponding to individuals with BLG were identified using the MaxQuant output files deposited in the related PRIDE repository (PXD008601; proteinGroups_workfile and allPeptides_reduced_from_proteinGroups), as the publication reported only the number of individuals in which milk proteins were identified. For studies published after 2021, we also recorded whether the Oral Signature Screening Database (OSSD) (Bleasdale et al., 2021) was applied to assess the preservation of the oral microbiome. As our goal was to evaluate enamel protein preservation in archaeological material, modern samples were excluded.

In total, the selected studies comprised 462 archaeological individuals, of which 434 had publicly available raw data. These corresponded to 498 raw files, as some individuals were represented by multiple files generated from different extraction methods (FASP, GASP, in-solution and SP3), different fractions (supernatant and pellet) and MS/MS analyses (Q-Exactive and Orbitrap Elite). The duplicates, triplicates, and quadruplicates were retained separately to assess potential differences across protocols and analytical replicates.

### 2.2 Bioinformatic analysis

The raw data from the 14 studies was publicly deposited on PRIDE and MassIVE repositories. Raw files from studies that had reanalysed datasets previously published by other authors were excluded (Hendy et al., 2018). All the raw files available (.raw) were re-analysed using MaxQuant version 2.4.6. and 2.6.7 (Cox and Mann, 2008) against a custom database comprising the commonly identified proteins of the human enamel proteome: Q99217| AMELX, Q99217-2| AMELX Isoform 2, Q99217-3| AMELX Isoform 3, Q99218| AMELY, Q99218-1|AMELY Isoform 1, Q9NRM1| enamelin (ENAM), Q9NP70| ameloblastin (AMBN), Q9NP70-2|AMBN Isoform 2, Q9NNX1 | tuftelin (TUFT1), Q9NNX1-2| TUFT1 Isoform 2, Q9NNX1-3| TUFT1 Isoform 3, Q6UX39| amelotin (AMTN), Q6UX39-2| AMTN Isoform 2, A1E959| odontogenic ameloblast associated protein (ODAM), O60882| matrix metalloproteinase-20 (MMP20), Q9Y5K2| kallikrein-4 (KLK4), Q9Y5K2-2| KLK4 Isoform 2 and Q9UMD9| Collagen alpha-1(XVII) (COL17A1). In addition, P53634| dipeptidyl peptidase (CATC) was included as a potential *in vivo* activator of KLK4 (Bartlett, 2013). The following non-specific proteins were also included: P02768| Serum albumin (ALBU), P02452| Collagen alpha-1(I) (CO1A1) and P08123| Collagen alpha-2(I) (CO1A2). A semi-specific search was conducted, with trypsin as the enzyme, and with Oxidation (M), Deamidation (NQ), Hydroxyproline and Phosphorylation (STY) as variable modifications.

MaxQuant outputs were analysed and visualised in R version 4.4.3 (R Core Team, 2025) using packages *tidyverse* version 2.2.0 (Wickham et al., 2019), *janitor* version 2.2.1 (Firke, 2024), *ggplot2* version 3.5.1 (Wickham, 2016), *dplyr* version 1.1.4 (Wickham et al., 2023) and *ggpubr* version 0.6.2 (Kassambara, 2025). Peptide counts were obtained from the Razor+Unique peptides reported in the MaxQuant proteinGroups output file and are summarized in Supplementary Information 1. Protein identifications were accepted when at least two Razor+Unique peptides were detected in a sample or replicate. For the calculation of the number of individuals identified with enamel proteins, the archaeological identifier was considered as the unique identifier, independently of the number of extraction identifiers available for a given individual. Moreover, cases with only one AMELX canonical peptide were further evaluated for AMELX isoform 3, which were included in the final counts of the number of individuals when ≥2 Razor+Unique peptides were observed. The same criteria was applied to AMBN individual counts when AMBN isoform 2 was detected in the absence of AMBN canonical peptides.

For statistical analyses, proteins were divided into two categories: (i) the total number of peptides recovered, which included enamel proteins (AMELX, AMELY, ENAM, AMBN, TUFT1, AMTN, ODAM, CATC, COL17A1, MMP20 and KLK4) together with human serum albumin (ALBU) and collagens (CO1A1 and CO1A2), and (ii) enamel proteins only.

The correlation between sample size (mg) and peptide counts (total number of proteins and enamel-only) was tested using Spearman’s rank correlation (ρ). Differences across extraction methods and peptide counts (total number of proteins and enamel-only) were assessed with Kruskal–Wallis non-parametric tests followed by pairwise Wilcoxon rank-sum tests with Benjamini–Hochberg correction (hereafter called pW-BH). Direct comparisons between FASP and SP3 were also performed as these were the two methods with the largest number of extractions (Supplementary Information). OSSD was evaluated against the total number of peptides recovered at the archaeological individual level. However, when an archaeological individual was represented by more than one extraction file (n=11), we excluded cases with discordant OSSD results across files (n=4) from further analysis. Dairy protein presence was evaluated against enamel peptides only and, similarly to the latter, we excluded cases with discordant dairy identifications (i.e., presence and absence across extractions; n=5). Sex estimation was assessed in archaeological individuals where osteological or genetic sex estimations were available and with AMELX and/or AMELY peptide sequences.

## 3. Results

### 3.1. Identification of the enamel proteome

The identification of enamel proteins per archaeological individual is summarised in Figure 1A and Table 1, and reported in the Supplementary Information. Proteins characteristic of the enamel proteome – AMELX, AMELY, AMBN, CATC, COL17A1, ENAM and MMP20 – were detected in ten out of fourteen studies (Figure 1B). Among these, AMELX was the most frequently identified protein (n=37 individuals), followed by AMBN (n=36 individuals), COL17A1 (n=28 individuals) and MMP20 (n=15 individuals). All other enamel proteins reported were detected in fewer than 10 individuals (Figure 1A), with the AMELY isoform 1 being reported in only three cases. Non-specific proteins, such as serum albumin and collagen type I (COL1A1 and COL1A2) were commonly identified, present in 89.6% (n=389) and 73.3% (n=318) individuals, respectively.

**Figure 1.**
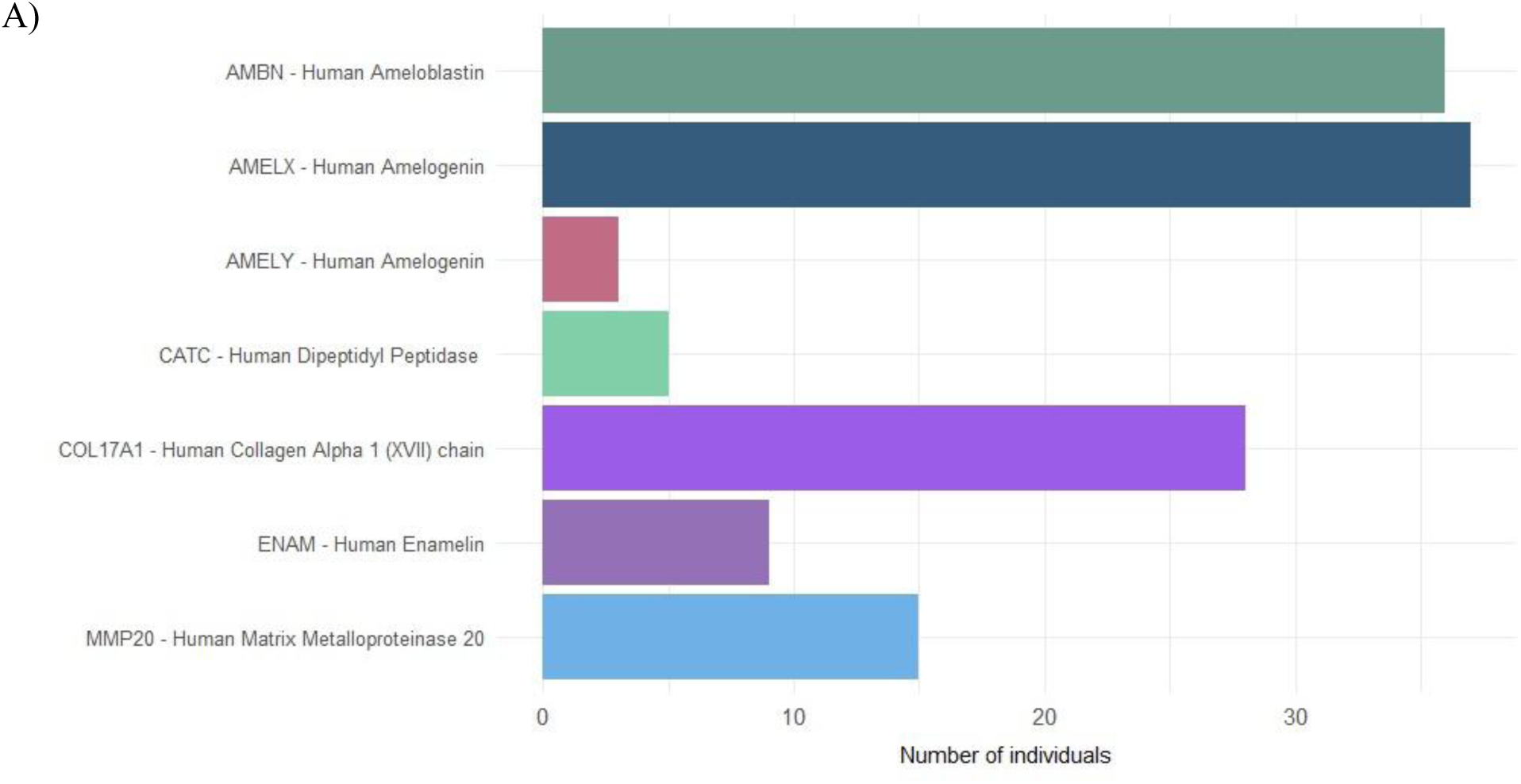
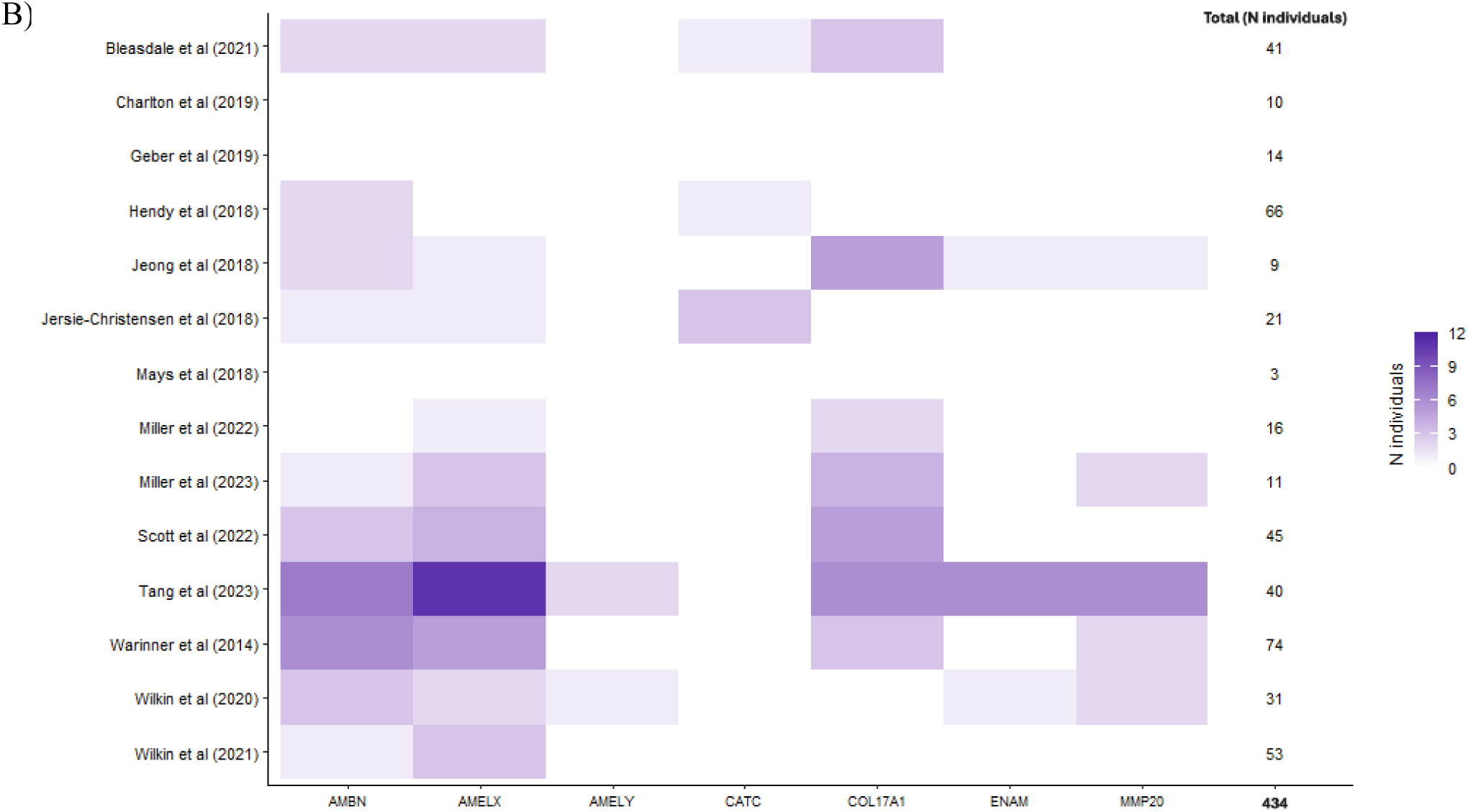
Enamel proteome identified (≥2 Razor + Unique peptides), in terms of A) Total of number of individuals in which they were present, out of a maximum of 434, and B) Number of individuals with enamel proteins identified, per study.

**Table 1.** Descriptive statistics of the enamel peptide identifications (≥2 Razor + Unique peptides) in archaeological individuals. AMELX and AMBN counts include the sum of AMELX canonical and AMELX isoform 3, and AMBN canonical and AMBN isoform 2, respectively.

| Protein Taxonomic Identification | N individuals | Razor + Unique peptides |  |  |  |
| --- | --- | --- | --- | --- | --- |
|  |  | Mean | SD | Min | Max |
| AMELX Human amelogenin | 37 | 6.5 | 5.7 | 2 | 26 |
| AMBN Human ameloblastin | 36 | 8.9 | 12.1 | 2 | 56 |
| COL17A1 Human collagen alpha 1(XVII) chain | 28 | 2.7 | 1.1 | 2 | 7 |
| MMP20 Human matrix metalloproteinase 20 | 15 | 5.9 | 6.1 | 2 | 23 |
| ENAM Human enamelin | 9 | 6.3 | 3 | 2 | 11 |
| CATC Human dipeptidyl peptidase | 5 | 2.2 | 0.4 | 2 | 3 |
| AMELY Human amelogenin isoform 1 | 3 | 2.3 | 0.6 | 2 | 3 |

### 3.2. Methodological, archaeological and dietary variables affecting peptide recovery

To evaluate the factors influencing enamel peptide recovery, methodological variables such as sample size, extraction methods and authentication through the OSSD were examined for total number of peptide counts (enamel and non-specific proteins, i.e., serum albumin and collagens) and enamel proteins only. Dietary variables were analysed exclusively for enamel proteins.

For the 176 samples where the starting material mass was reported, values ranged from 1.0 to 145.4 mg. Within this range, enamel proteins were identified in samples weighing between 5.0 and 32.9 mg. No correlation was observed between sample mass and the total number of dental enamel peptides, or between sample mass and peptides from enamel-specific proteins (Spearman’s rank correlation, sample mass∼total number of peptides ρ=-0.04 and *p*=0.57; sample mass∼total number of enamel peptides ρ=-0.06 and *p*=0.44), indicating that the amount of starting material did not influence dental enamel peptide recovery. When considering the extraction protocol selected (n=498 samples), significant differences were identified both for the total number of peptides (Kruskal–Wallis χ^²^=42.09, *p*=3.84×10^⁻⁹^) and for enamel-specific peptides (Kruskal–Wallis χ^²^=32.995, *p*=3.23×10^⁻7^). Pairwise comparisons showed that GASP yielded significantly fewer total number of peptides (enamel and non-specific) than FASP, SP3, and in-solution digestion (pW-BH *p*<0.05 in all cases), while SP3 and in-solution digestion yielded higher median peptide counts than FASP (pW-BH *p*<0.05 in both cases, Figure 2) but did not differ from each other. However, as the in-solution digestion method was only applied to a small number of samples (n=22, Figure 2), this represents an unbalanced comparison. We therefore focused on the two most widely used methods for dental calculus protein extraction published thus far, FASP and SP3, to assess which method yielded a higher count of enamel and non-specific peptides.

**Figure 2.**
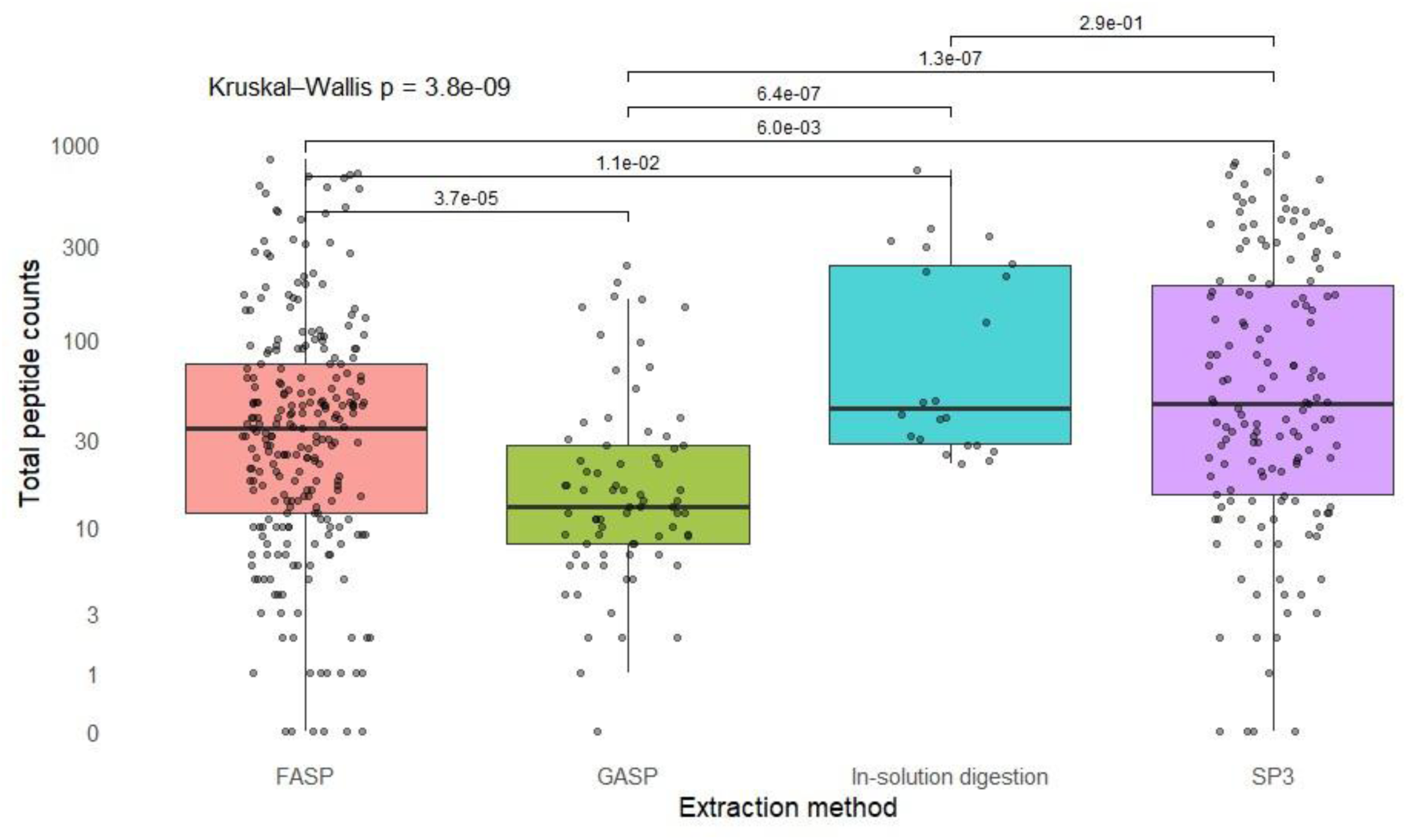
Distribution of total peptide identifications (≥2 Razor+Unique peptides) in ancient dental calculus by extraction method (FASP, GASP, In-solution digestion and SP3). P-values from pairwise Wilcoxon rank-sum tests with Benjamini–Hochberg correction are shown. Each individual point represents an extract.

For total peptide recovery, both FASP and SP3 yielded successful identifications in the majority of the extractions (FASP=95% and SP3=97%). However, SP3 yielded higher peptide counts than FASP (Table 2), a trend also observed in paired comparisons of individuals extracted by both methods, where SP3 outperformed FASP in 8 of 11 cases. When restricted to enamel-specific peptides, mean peptide counts were slightly higher for SP3 (20.5±30.6) than FASP (10.4±14.6), with enamel proteins identified in 28 and 33 extractions for SP3 and FASP, respectively. Nevertheless, enamel peptides were recovered more frequently with SP3 (19%) than with FASP (13%), with higher maximum yields (FASP max=66; SP3 max=122).

**Table 2.** Descriptive statistics of the number of peptide identifications (≥2 Razor+Unique peptides) identified in ancient human calculus samples, reported separately for total peptides (enamel and non-specific proteins) and for enamel-only peptides.

| Method | N tested | Total number of peptides |  |  |  |  | Total number of enamel peptides |  |  |  |  |
| --- | --- | --- | --- | --- | --- | --- | --- | --- | --- | --- | --- |
|  |  | N identified | Median | Mean | SD | Max | N identified | Median | Mean | SD | Max |
| FASP | 253 | 240 | 36 | 84.9 | 141.2 | 825 | 33 | 5 | 10.4 | 14.6 | 66 |
| SP3 | 149 | 144 | 47 | 146.3 | 193.3 | 870 | 28 | 6 | 20.5 | 30.6 | 122 |

To determine whether other analytical parameters influenced peptide recovery, we examined extraction fractions. In the three archaeological individuals from Stonehenge (Mays et al., 2018) where both supernatant and pellet fractions were analysed with GASP, only non-specific proteins were identified. Supernatant fractions yielded significantly higher peptide counts than pellets (Wilcoxon rank-sum test *p*=0.008; n=6), whereas no significant differences were observed between LC-MS/MS run times (1h vs 3h; Wilcoxon rank-sum test *p*>0.05; n=6).

Following this, we examined whether authentication through the application of OSSD corresponded to dental enamel peptide recovery. Of the 157 individuals for which OSSD validation was applied, 109 passed and 48 failed the threshold, with the latter excluded from subsequent analyses. Despite this, both groups yielded comparable total peptide counts (mean failure=170±231 peptides; mean pass=114±162 peptides, Wilcoxon rank-sum test *p*=0.63). Dental enamel peptides were identified in a small subset of both OSSD-failed (AMELX and AMELY present in seven and one individuals, respectively) and OSSD-passed samples (AMELX present in fourteen individuals and AMELY present in one, respectively).

We also compared dental enamel peptide occurrence with the recovery of dairy proteins. Among the 429 individuals analysed for dairy proteins, 187 showed evidence of milk consumption and 242 did not yield identifiable dairy proteins based on the original publications. Dental enamel peptides were identified in both groups, with AMELX detected in 12 dairy-positive and 25 dairy-negative individuals. Among the individuals with AMELX peptides, AMELY was identified in 2 dairy-positive individuals and 1 dairy-negative individual. No significant difference was observed in dental enamel peptide counts between individuals with and without evidence of dairy consumption (Wilcoxon rank-sum test *p*=0.72; n=63 individuals). These findings show that enamel proteins can be recovered both in individuals with and without dairy proteins, highlighting their complementarity in multidisciplinary approaches. Specifically, the detection of amelogenin peptides alongside the identification of dairy proteins offers an additional line of evidence for possible sex estimation.

### 3.3 Application to sex estimation

Overall, amelogenin peptides were identified in 37 individuals, of which nine had been previously assigned a biological sex through genetic or osteological methods (Table 3). In four of these nine cases, the proteomic results were consistent with previous results, that is, females showed only AMELX peptides. However, two genetically and osteologically identified males, as well as one osteologically identified male, exhibited only AMELX peptide signals. The absence of AMELY peptides in these individuals may reflect differential preservation or lower peptide abundance of AMELY relative to AMELX, as only ∼10% of amelogenin transcripts in males derive from Y-chromosome, whereas >90% originate from X-chromosome (Salido et al., 1992; Stewart et al., 2016; Yamakoshi, 2014). In contrast, the Q99218-1| AMELY-(58-64) peptide (WYQSM(ox)IRPPYS) was detected in one individual identified as female based on osteological methods.

**Table 3.** Biological sex estimation based on osteological and genetic methods, and corresponding amelogenin peptide counts (≥2 Razor+Unique peptides) per individual, grouped by study. NA – Not available.

| Study | Archaeological ID | Osteological Sex | Genetic Sex | Q99217\|<br>AMELX | Q99217-3\|<br>AMELX-3 | Q99218-1\|<br>AMELY-1 |
| --- | --- | --- | --- | --- | --- | --- |
| Warinner et al (2014) | KAL930 | NA | NA | 6 | 1 | 0 |
| Warinner et al (2014) | KAL934 | NA | NA | 6 | 1 | 0 |
| Warinner et al (2014) | STH276 | NA | NA | 2 | 0 | 0 |
| Warinner et al (2014) | STH419 | NA | NA | 2 | 1 | 0 |
| Warinner et al (2014) | STH421 | NA | NA | 2 | 1 | 0 |
| Jeong et al (2018) | 2007_9 | Male | Male | 17 | 2 | 1 |
| Jeong et al (2018) | 2007_14 | Female | Female | 23 | 3 | 0 |
| Jersie-Christensen et al (2018) | #13 A1635 | Male | NA | 6 | 0 | 0 |
| Wilkin et al (2020) | AT-233 | NA | NA | 9 | 0 | 0 |
| Wilkin et al (2020) | AT-360 | NA | NA | 17 | 4 | 2 |
| Wilkin et al (2020) | AT-650 | NA | NA | 9 | 1 | 0 |
| Bleasdale et al (2021) | Box 32G | NA | NA | 2 | 1 | 0 |
| Bleasdale et al (2021) | Trench IV | NA | NA | 3 | 0 | 0 |
| Wilkin et al (2021) | MUR2 N-130.1 | NA | NA | 1 | 2 | 0 |
| Wilkin et al (2021) | MUS5 N-1 | NA | NA | 2 | 0 | 0 |
| Wilkin et al (2021) | UTE6 K.4 N-5 | NA | NA | 2 | 1 | 0 |
| Miller et al (2022) | DA-AKH-16 | NA | NA | 4 | 1 | 0 |
| Miller et al (2023) | HOR-01 | NA | NA | 2 | 0 | 0 |
| Miller et al (2023) | HOR-27 | NA | NA | 6 | 0 | 0 |
| Miller et al (2023) | HOR-48 | NA | NA | 4 | 2 | 0 |
| Scott et al (2022) | SGR002 | NA | NA | 8 | 3 | 0 |
| Scott et al (2022) | VSK002 | NA | Female | 4 | 1 | 0 |
| Scott et al (2022) | KSI001 | NA | Female | 9 | 3 | 0 |
| Scott et al (2022) | KBD001 | Male | Male | 3 | 2 | 0 |
| Tang et al (2023) | 2003LWZLJM6 | NA | NA | 1 | 3 | 0 |
| Tang et al (2023) | 2017LZTCM1-60 | NA | NA | 7 | 21 | 3 |
| Tang et al (2023) | 2017ZSM7 | NA | NA | 3 | 0 | 0 |
| Tang et al (2023) | 2018ZGM1 | NA | NA | 1 | 8 | 0 |
| Tang et al (2023) | 2018ZSEM3 | Female | NA | 26 | 6 | 1 |
| Tang et al (2023) | 2018ZSM5 | NA | NA | 4 | 0 | 0 |
| Tang et al (2023) | 2019BZM2 | NA | NA | 8 | 11 | 0 |
| Tang et al (2023) | 2019QLSZM1 | Female | NA | 11 | 2 | 2 |
| Tang et al (2023) | 2019RRNCG | NA | NA | 5 | 8 | 0 |
| Tang et al (2023) | 2019ZGM7 | NA | NA | 2 | 0 | 0 |
| Tang et al (2023) | 2019ZMM1 | Female | NA | 9 | 1 | 0 |
| Tang et al (2023) | 2019ZSEM25 | NA | NA | 10 | 12 | 0 |
| Tang et al (2023) | 2019ZSEM31 | NA | NA | 1 | 2 | 0 |

Interestingly, two previously undetermined individuals also yielded unique AMELY peptide signals: one yielded two unique sequences, AMELY(46-54) and AMELY-(58-64), while the other one exhibited three, AMELY(46-54), AMELY-(58-64) and AMELY-(110-122). These results suggest that amelogenin identified in ancient calculus may provide information for biological sex estimation, especially in otherwise indeterminate cases.

## 4. Discussion

Proteins have the potential to remain well-preserved within the highly mineralized structure of dental enamel, providing valuable insights into taxonomic identification, phylogenetic relationships and biological sex determination (Gil-Bona and Bidlack, 2020; Mikšík et al., 2023; Taurozzi et al., 2024). Until now, however, the dental enamel proteome has only been detected within enamel tissue itself. This study broadens that scope by demonstrating that dental enamel-specific proteins can also be identified in ancient dental calculus, thereby introducing a new line of research in a matrix already recognised as a rich source of biomolecular evidence (Fagernäs and Warinner, 2023). Understanding the processes occurring on the enamel surface during plaque biofilm formation and mineralisation is crucial for interpreting how enamel proteins become incorporated and preserved.

### 4.1. Detection of enamel proteins in ancient calculus

The tooth enamel surface is continuously subjected to demineralization and remineralization cycles throughout life (Enax et al., 2024). When dietary fermentable carbohydrates are metabolised by cariogenic bacteria, organic acids are produced causing a rapid decrease in pH (Featherstone, 2008; García-Godoy and Hicks, 2008; Pitts et al., 2017). Once the pH reaches the critical threshold for enamel (approximately pH 5.5), hydroxyapatite begins to dissolve, releasing calcium and phosphate ions from the tooth surface into the surrounding oral environment (Featherstone, 2008, 2004; Koontongkaew et al., 2024). Tenuta et al. (2006) demonstrated that following this acid-induced demineralisation of dental enamel, calcium and phosphate concentrations within the plaque biofilm increase markedly before returning to baseline values after around 24 hours. This temporary window of ionic exchange allows enamel-derived ions to interact with the developing plaque biofilm (Enax et al., 2024; Tenuta et al., 2006). Repeated acidic episodes driven by continued acid challenge and pH drops disrupt the demineralization-remineralization balance, resulting in a net loss of dental tissue and the formation of early carious lesions (Kidd and Fejerskov, 2004; Koontongkaew et al., 2024; Pitts et al., 2017). Although dental enamel protein release during these events has not been experimentally demonstrated, we hypothesize that the same cycles of tooth surface dissolution that mobilise mineral and non-mineral ions may also release dental enamel protein fragments, which could then be entrapped by the mineralising plaque matrix. As calculus accumulates through repeated cycles of plaque formation and mineralization, these episodic dissolution events at the dental enamel surface may therefore facilitate the incorporation of dental enamel-derived proteins.

A second and non-exclusive possibility relates to mechanical abrasion of the tooth surface during archaeological dental calculus sampling. The application of excessive force on specimens where calculus is strongly adhered to the tooth surface may cause minor enamel scraping, inadvertently introducing enamel proteins into the sample. Although this risk is low when standard procedures are followed (e.g., Sabin and Yates, 2020), it remains a factor that must be considered when interpreting enamel protein presence in calculus.

Amelogenin was the most frequently detected enamel protein in our study, followed by ameloblastin, while enamelin was identified only in a small subset of individuals. This pattern closely aligns with previous enamel proteome studies, in which lower amounts of ameloblastin and enamelin are expected as their expression ceases at the early maturation stage (Castiblanco et al., 2015; Stewart et al., 2016). In addition to these structural proteins, we also detected other enamel peptides such as MMP20 and CATC, although KLK4 itself was not identified in this study. We further recovered COL17A1, a protein involved in ameloblast attachment to the developing enamel surface (Asaka et al., 2009), in frequencies comparable to those of ameloblastin.

Non-specific proteins, including serum albumin and collagen type I, were frequently recovered. The detection of serum albumin, one of the bulk salivary proteins, is expected given that the plaque biofilm is continuously exposed to saliva and gingival crevicular fluid in the oral cavity (Hu et al., 2005; Jin and Yip, 2002). Albumin has also been suggested to enter teeth *in vivo* via porous dental enamel or as a component of the non-amelogenin fraction during enamel development (Gil-Bona and Bidlack, 2020). On the other hand, the presence of collagen peptides may derive from multiple sources, including handling, laboratory contamination or inadvertent co-extraction of bone powder when calculus is removed from teeth *in situ*. Mackie and colleagues (2017) explored whether the presence of collagen could be the result of alveolar bone resorption due to periodontitis but found no relationship between skeletal lesions of periodontal disease and collagen detection, suggesting that its presence is possibly related to contamination during sampling. Although contamination could not be ruled out, further studies exploring the incorporation of collagen in calculus in living populations may help us understand whether its presence could be associated with underlying oral pathologies.

### 4.2. Factors affecting preservation of the enamel proteome

The preservation of dental enamel proteins in ancient calculus suggests that recovery might be influenced primarily by extraction method and site-specific preservation conditions. The highest protein yields were obtained in studies that applied FASP and SP3, the two most widely used methods in dental calculus palaeoproteomics. In contrast, GASP and in-solution digestion tended to produce lower recoveries, likely because GASP requires substantial clean-up to isolate the proteins from the gel matrix, increasing the risk of protein loss, while direct solubilisation and digestion approaches lack additional clean-up steps and can therefore also lead to reduced peptide recovery (Cleland, 2018; Palmer et al., 2021). When comparing the best-performing protocols, SP3 showed higher recovery of dental enamel peptides. This may reflect the fact that, unlike FASP, SP3 does not rely on molecular weight and, due to the combination of both hydrophobic and hydrophilic magnetic beads, can capture a broader range of protein fragments, which is advantageous in the context of poorly preserved archaeological material (Cleland, 2018). It should also be noted that all the extracts analysed here were digested with trypsin. As the enamel proteome is digested *in vivo* (Bartlett et al. 2013), this additional digestion may lead to peptides that are too short to be analysed by mass spectrometry, and the results presented here may therefore be an underestimation of the amount of enamel proteins that are integrated into dental calculus.

Although dental enamel is the hardest and most degradation-resistant tissue in the human skeleton, dental enamel peptides incorporated into dental calculus might be exposed to a distinct microenvironment within the plaque matrix and are therefore subjected to different preservation dynamics than those preserved within intact enamel. While dental calculus is a robust reservoir of diverse biomolecules, it is, in comparison to dental enamel, more susceptible to post-depositional alteration. Burial conditions such as soil chemistry, humidity, temperature and pH, as well as different burial practices, may influence the degree of preservation of proteins harboured in dental calculus (Mackie et al., 2017b; Pedergnana et al., 2025). Further research into the depositional environments of the archaeological sites analysed here would shed light on how site-specific taphonomic conditions may have influenced the survival of enamel-derived peptides. Moreover, previous studies (e.g., Fagernäs et al., 2022; Mackie et al., 2017b) have shown individual variability in both protein and DNA preservation in calculus, which may relate to the location of tooth sampled (anterior vs posterior dentition), size of the dental calculus deposit, differences in oral health, or *in vivo* processes occurring within the oral cavity as, for example, episodic events of acid-induced demineralisation of dental enamel. The intra-individual variability of the oral environment, combined with post-depositional conditions, are factors that should be considered when interpreting the differences found between individuals even within archaeological sites.

### 4.3. Potential as a tool for sex estimation

Biological sex estimation via traditional osteological methods poses several challenges, as it is highly dependent on skeletal preservation and completeness (Díaz-Zorita Bonilla et al., 2024; Mikšík et al., 2023). In archaeological assemblages, the skull and *os coxae* – the preferred skeletal elements for analysis – are often heavily fragmented or absent, reducing the accuracy of osteological estimates (Bružek et al., 2024; Godinho et al., 2025). Moreover, complex funerary behaviours involving remobilisation and potential secondary depositions often result in the dissociation of skeletal elements of the individuals (Evangelista and Godinho, 2020; Filipe et al., 2013; Godinho, 2008; Godinho et al., 2019; Valera et al., 2026, 2018), thus precluding hip bone based morphological sex estimation. Consequently, biomolecular methods have emerged as reliable and alternative methods for sex determination.

We have shown that AMELX and AMELY sex-specific peptide sequences can be identified in ancient dental calculus. Detection of AMELY in three archaeological cases suggests male individuals, while its absence in other cases could result from either a low abundance of AMELY-specific peptides, or a female sex. Similarly to proteomic sex estimation using tooth enamel, this approach has the potential to be applied to incomplete individuals or even isolated teeth. Of the 37 individuals from whom amelogenin peptides were recovered, we were able to compare our results with genomic data in six cases and with osteological estimation in seven. Within this subset, four individuals did not match previously reported genomic or osteological estimates. Buonasera et al. (2020) compared biological sex estimation through proteomics, genomics and osteological methods, reporting accuracies of 100%, 91% and 51%, respectively. The agreement among methods was high, however conflicts were reported. Future integration of these dental calculus proteomic results with osteological and genomic data will be crucial for clarifying the cases where the sex estimation does not match and to further assess the reliability of sex estimation using proteins derived from calculus.

Palaeoproteomic data generated from archaeological specimens is usually accessible and reusable due to the increased adherence to open data-sharing practices (Chiang et al., 2024; Dekker et al., 2025). This is particularly relevant as archaeological material is a finite resource and often cannot be resampled, especially in the case of dental calculus that, although ubiquitous, varies greatly in the quantity available for analysis (Fagernäs and Warinner, 2023). The growing use of palaeoproteomics applied to calculus has also resulted in a marked increase in sampling requests to museums and heritage institutions, at times employing sampling strategies that include the complete removal of the calculus deposits or the pooling of material across several individuals (Gancz et al., 2024). Our study showcases the importance of re-using existing datasets to test new hypotheses and to further advance our knowledge in the field. Reanalysis of previously generated data not only eliminates the need for destructive sampling of new archaeological specimens but also demonstrates that new research questions and approaches can uncover biomolecules preserved in the dental calculus matrix that had not been reported in earlier studies.

## Conclusions

This study shows that enamel proteins can be identified in ancient dental calculus, expanding the range of biomolecules known to be preserved in this matrix. The detection of amelogenin, ameloblastin, enamelin, MMP20, COL17A1 and CATC indicates that, under certain conditions, proteins characteristic of the enamel proteome may enter the plaque matrix. Recovery patterns across archaeological sites suggest that extraction methods and site-specific preservation conditions have a greater effect on peptide detection. The identification of AMELX and AMELY peptides in calculus also demonstrates that enamel proteins preserved in this matrix can possibly contribute to sex estimation in previously undetermined archaeological cases. However, discrepancies with genomic and osteological assessments highlight the need for integrated approaches.

Palaeoproteomics is growing at a rapid pace in recent years, with a greater number of archaeological materials yielding proteins that shed light on long-debated questions in archaeology, bioarchaeology, paleoanthropology or zooarchaeology. Despite the significance of these discoveries, there is an increasing concern regarding the destructive nature of sampling as archaeological samples are unique and irreplaceable. Progress in the field should also rely on systematic reanalysis of existing datasets to explore new hypotheses without requiring the sampling of new archaeological material. Our study demonstrates that when raw data are made available, re-analysis can be extremely valuable and can lead to the identification of proteins not reported in the original studies.

## Funding

A.L. is supported by the Portuguese Foundation for Science and Technology (FCT; Ph.D. Grant No. 2022.13780.BD; DOI: https://doi.org/10.54499/2022.13780.BD) and the Archaeology of Portugal Fellowship from the Archaeological Institute of America. This research has been made possible through funding from the European Research Council (ERC) under the European Union’s Horizon 2020 research and innovation programme, grant agreement no. 948365 (PROSPER, awarded to F.W.), the European Union’s Horizon Europe research and innovation programme under the Marie Skłodowska-Curie grant agreement no. 101106627 (PROMISE, awarded to Z.F.) and no. 101208619 (ProPGen, awarded to V.V.I). R.M.G. is funded by Fundação para a Ciência e a Tecnologia (FCT; contract reference 2023.10993.TENURE.006). This research was also supported by the FCT R&D research project “ParaFunction” (project reference 2022.07737.PTDC; https://doi.org/10.54499/2022.07737.PTDC).

## Acknowledgements

The authors acknowledge ICArEHB – Interdisciplinary Center for Archaeology and Evolution of Human Behaviour, funded by the Portuguese Foundation for Science and Technology (FCT) under program UID/04211/2025. A.L. acknowledges the Archaeological Association of the Algarve. For the purpose of Open Access, the authors have applied a CC-BY public copyright license to any Author‘s Accepted Manuscript (AAM) version arising from this submission.

## Author contributions

**Conceptualization:** A.L., F.W., Z.F.; **Data analysis**: A.L., V.V.I., Z.F. with input from F.W.; **Writing-original draft preparation**: A.L., Z.F.; **Writing-review and editing**: A.L.; F.W., R.E.G., R.M.G, V.V.I, Z.F.

## Data availability

The research data is available as supplementary material and R code used for analysis is archived in Zenodo at https://doi.org/10.5281/zenodo.19135248

## Notes

https://zenodo.org/records/19135248

### Competing Interest Statement

The authors have declared no competing interest.

